# Replication stress marker phospho-RPA2 predicts response to platinum and PARP inhibitors in homologous recombination-proficient ovarian cancer

**DOI:** 10.1101/2024.11.21.624682

**Authors:** Angela Schab, Amanda Compadre, Rikki Drexler, Maggie Loeb, Kevin Rodriguez, Joshua Brill, Shariska Harrington, Carmen Sandoval, Brooke Sanders, Lindsay Kuroki, Carolyn McCourt, Andrea R. Hagemann, Premal Thaker, David Mutch, Matthew Powell, Violeta Serra, Ian S. Hagemann, Ann E. Walts, Beth Y. Karlan, Sandra Orsulic, Katherine Fuh, Lulu Sun, Priyanka Verma, Elena Lomonosova, Peinan Zhao, Dineo Khabele, Mary Mullen

## Abstract

**Background:** Ovarian cancer treatment includes cytoreductive surgery, platinum-based chemotherapy, and often poly (ADP-ribose) polymerase (PARP) inhibitors. Homologous recombination (HR)-deficiency is a well-established predictor of therapy sensitivity. However, over 50% of HR-proficient tumors also exhibit sensitivity to standard-of-care treatments. Currently, there are no biomarkers to identify which HR-proficient tumors will be sensitive to standard-of-care therapy. Replication stress may serve as a key determinant of response.

**Methods:** We evaluated phospho-RPA2-T21 (pRPA2) foci via immunofluorescence as a potential biomarker of replication stress in formalin-fixed, paraffin-embedded tumor samples collected at diagnosis from patients treated with platinum chemotherapy (discovery cohort: n = 31, validation cohort: n = 244) or PARP inhibitors (n = 87). Recurrent tumors (n = 37) were also analyzed. pRPA2 scores were calculated using automated imaging analysis. Samples were defined as pRPA2-High if > 16% of cells had ≥ 2 pRPA2 foci.

**Results:** In the discovery cohort, HR-proficient, pRPA2-High tumors demonstrated significantly higher rates of pathologic complete response to platinum chemotherapy than HR-proficient, pRPA2-Low tumors. In the validation cohort, patients with HR-proficient, pRPA2-High tumors had significantly longer survival after platinum treatment than those with HR-proficient, pRPA2-Low tumors. Additionally, the pRPA2 assay effectively predicted survival outcomes in patients treated with PARP inhibitors and in recurrent tumor samples.

**Conclusion:** Our study underscores the importance of considering replication stress markers alongside HR status in therapeutic planning. Our work suggest that this assay could be used throughout a patient’s treatment course to expand the number of patients receiving effective therapy while reducing unnecessary toxicity.

## INTRODUCTION

Ovarian cancer is treated with a combination of cytoreductive surgery and platinum-based chemotherapy (1-4). Additionally, current guidelines offer poly (ADP-ribose) polymerase (PARP) inhibitors, such as olaparib or niraparib, as maintenance therapy for all patients who are PARP inhibitor naïve and have had a partial or complete response to platinum chemotherapy (5-7). While these therapies are effective initially, over 85% of patients with advanced ovarian cancer eventually develop tumor resistance to standard-of-care platinum chemotherapy and PARP inhibitors and die within five years of diagnosis (1-4). A major determinant of tumor sensitivity or resistance to these therapies is whether the tumor can perform homologous recombination (HR) to repair double-stranded DNA (dsDNA) breaks created by platinum-induced DNA adducts or PARP inhibitors (8). Previously, we developed an automated immunofluorescence assay to evaluate functional HR in formalin-fixed, paraffin-embedded (FFPE) ovarian tumors (9). Our assay measures foci of RAD51, as binding of this protein to DNA relies on many upstream HR events and is a functional read-out for HR proficiency (10, 11). We and others found that while most RAD51-Low or functionally HR-deficient tumors respond to therapy, over 50% of RAD51-High or HR-proficient tumors are also sensitive to these therapies. This suggests that HR-independent mechanisms play a significant role in determining therapeutic response (9, 12, 13). These findings are supported by clinical trial data demonstrating that over 65% of women with BRCA-mutated or HR-deficient tumors achieve a survival rate exceeding 90 months when treated with platinum chemotherapy followed by PARP inhibitors (9, 12-14). Similarly, there is a subset of up to 30% of women with HR-proficient tumors that experience survival extending beyond 70 months after platinum and PARP inhibitor treatment (15, 16). This is in stark contrast to the median survival for women with HR-proficient tumors after platinum and PARP inhibitor treatment of 36.6 months (16). Together, these data indicate the need for a more accurate assay to determine whether patients with HR-proficient tumors will benefit from platinum chemotherapy and PARP inhibitors. Such an assay could improve treatment decisions, expand the number of patients receiving effective therapy, and reduce unnecessary toxicity (17-22).

An important determinant of platinum and PARP inhibitor sensitivity in HR-proficient tumor cells *in vitro* is replication stress (23, 24). Replication stress occurs when, in response to genotoxic damage, replication forks slow and initiate DNA damage tolerance pathways leading to the accumulation of single-stranded DNA (25, 26). If the cell repairs the damage, then the replication stress resolves. However, if replication stress is excessive, the replication fork collapses, resulting in multiple double-stranded DNA breaks and cell death even in HR-proficient cells. Specifically, replication stress events such as fork degradation and single-stranded DNA (ssDNA) gap formation, are directly linked to treatment sensitivity independent of HR-status (23, 24, 27-29). A key protein involved in replication stress is replication protein A (RPA). RPA, which is composed of three subunits, binds to the single-stranded DNA that accumulates in response to stalled replication forks. Next, one of the three subunits, RPA2, is phosphorylated sequentially at multiple sites. Thr21 is the last site to be phosphorylated and indicates impending replication catastrophe and irreparable damage (30-32). Therefore, we hypothesized that HR-proficient tumor cells with high Thr21 phospho-RPA2 (pRPA2) would have a better response to platinum chemotherapy or PARP inhibitors than RAD51-High tumors with low pRPA2.

Here, we show that pRPA2 foci can be reliably identified in FFPE tumor samples. Additionally, using samples from multiple cohorts of patients with ovarian cancer, we show that HR-proficient, pRPA2-High tumors had significantly better responses to platinum chemotherapy and PARP inhibitors than HR-proficient, pRPA2-Low tumors. We also developed automated quantification methods to objectively measure functional HR status and pRPA2 foci, which should enable testing of this new assay in clinical trials.

## RESULTS

### pRPA2 foci reflect replication stress and can be quantified in patient-derived ovarian cancer samples

We hypothesized that HR-proficient tumor cells with high Thr21 phosphorylated RPA2 (pRPA2) foci would respond better to platinum chemotherapy than those with low pRPA2 foci. We first stained nine genomically HR-proficient primary ovarian cancer cells (POVs) (Supplementary Figure 1A) with an antibody against pRPA2 and counted the number of pRPA2 foci at baseline per cell. We then calculated *in vitro* carboplatin sensitivity (IC_50_) for each POV. The number of pRPA2 foci significantly negatively correlated with carboplatin IC_50_ (Pearson r= -0.90, *P*<0.001) (Supplementary Figure 1B). These data suggested that pRPA2 is a useful biomarker for carboplatin sensitivity in patient-derived ovarian cancer cells.

Non-fixed primary cells are not readily available in the clinic. Therefore, to facilitate the translation of our findings, we next sought to determine the accuracy of pRPA2 as a biomarker for replication stress in FFPE samples. To do so, we treated two HR-proficient high-grade serous ovarian cancer cell lines (TYKNU and OVCAR8) with increasing doses of hydroxyurea to induce replication stress. We then formalin-fixed and paraffin-embedded the lines as cell blocks and evaluated pRPA2 foci by immunofluorescence (9). Each sample was assigned a pRPA2 score, defined as the percentage of cells with ≥ 5 pRPA2 foci. With increasing doses of hydroxyurea, we observed increasing pRPA2 scores (Supplementary Figure 2A-B). To validate the specificity of pRPA2 Thr21 antibody, we used siRNA to knock down RPA2 in the same cell lines, created FFPE cell blocks, and evaluated pRPA2 foci. We observed significantly lower pRPA2 scores in cells in which RPA2 was knocked down, both with and without hydroxyurea treatment (Supplementary Figure 2C-D). We conclude that the pRPA2 Thr21 antibody is sensitive and specific, that pRPA2 foci staining accurately reflects replication stress, and that the assay can be conducted on FFPE tumor samples.

### pRPA2 foci in FFPE patient samples predict response to platinum chemotherapy and survival outcomes in RAD51-High tumors

To investigate the accuracy of pRPA2 scores in predicting response to platinum chemotherapy, we evaluated the association between pRPA2 foci and platinum chemotherapy response in a discovery cohort of biopsies obtained from primary tumor sites before patients received neoadjuvant carboplatin (n=31) (Table 1). All patients had advanced-stage disease and at least a component of high-grade serous histology. All samples were screened by a board-certified pathologist for diagnosis, cellularity, and necrosis. Samples had previously been scored as RAD51-High (HR-proficient) or RAD51-Low (HR-deficient) using RAD51 foci as described (9).

**Table 1:** Patient Characteristics.

|  | Discovery Cohort (n=31) | Validation Cohort Platinum (n=244) | Validation Cohort PARP Inhibitor (n=87) | Validation Cohort Recurrent Tumor (n= 38) |
| --- | --- | --- | --- | --- |
| Age (years) | 67.1 ± 10.1 | 62.7 ± 11.9 | 60.7 ± 11.3 | 49.2 ± 19.8 |
| Race |  |  |  |  |
| Black | 6 (19.4) | - | - | 2 (5.1) |
| White | 25 (80.6) |  |  | 32 (82.1) |
| Other | 0 (0) |  |  | 5 (12.8) |
| FIGO Stage |  |  |  |  |
| I | 0 | 2 (0.8) | 1 (1.1) |  |
| II | 0 | 11 (4.5) | 5 (5.8) | 2 (5.3) |
| III | 25 (80.6) | 183 (75.0) | 60 (69.0) | 30 (78.9) |
| IV | 6 (19.4) | 48 (19.7) | 21 (24.1) | 6 (15.8) |
| BRCA Mutation** |  |  |  |  |
| None | 21 (67.7) | 107 (43.9) | 49 (56.3) | 13 (34.2) |
| BRCA1 | 6 (19.4) | 59 (24.2) | 26 (29.9) | 7 (18.4) |
| BRCA2 | 1 (3.2) | 26 (10.6) | 11 (12.6) | 2 (5.3) |
| No testing/Missing | 3 (9.7) | 52 (21.3) | 1 (1.2) | 16 (42.1) |
| Chemotherapy Response Score (CRS) |  |  |  |  |
| 1-2 | 18 (58.1) | - | - | - |
| 3 (pCR) | 13 (41.9) |  |  |  |
| Neoadjuvant Chemotherapy |  |  |  |  |
| Yes | 31 (100) | 50 (20.5) | 32 (36.8) | 0 |
| No | 0 | 150 (61.5) | 52 (59.8) | 38 (100) |
| Missing | 0 | 44 (18.0) | 3 (3.4) | 0 |
| Platinum Sensitivity |  |  |  |  |
| Sensitive (PFI≥ 6m) | 25 (80.6) | 212 (86.9) | 84 (96.6) | 37 (97.4) |
| Resistant (PFI< 6m) | 6 (19.4) | 27 (11.1) | 3 (3.4) | 1 (2.6) |
| Missing | 0 | 5 (2.0) | 0 | 0 |
| Residual Disease at Cytoreduction |  |  |  |  |
| Absent | 22 (71.0) | 156 (63.9) | 7 (8.0) | - |
| Present | 7 (22.6) | 25 (10.3) | 76 (87.4) |  |
| Missing | 2 (6.4) | 63 (25.8) | 4 (4.6) |  |
| Median Follow-up (months) | 41.9 ± 22.7 (1.1 - 88.7) | 59.6 ± 41.6 (5.0 – 276.0) | 54.7 ± 31.1 (12.9-173.4) | 66.7 ± 36.6 (15.8-156.5) |

Data are n (%) unless stated otherwise. ± Denotes standard deviation. PFI, Progression free interval. \*\*BRCA mutation status determined by germline testing or whole genome sequencing. Limited information regarding pathogenicity of mutations.

All tumors had contributive results. pRPA2 foci were manually counted in all tumors, and all samples had valid, quantifiable pRPA2 foci (0-31 per cell). Two dichotomization cutoffs – the number of pRPA2 foci per cell and the percentage of positive cells – were defined by optimizing the progression-free survival (PFS) difference in HR-proficient tumors (33). Tumors were defined as pRPA2-High if >20% of cells had ≥2 pRPA2 foci and pRPA2-Low if ≤20% of cells had ≥2 pRPA2 foci. Eight (25.8%) tumors were pRPA2-High, and 23 (74.2%) were pRPA2-Low (Figure 1A, B). In the analysis in which RAD51 status was not considered, patients with pRPA2-High tumors had similar PFS (17.1 vs 9.9 months, *P*=0.4) and overall survival (OS) (47.1 vs 32.7 months *P*=0.3) as those with pRPA2-Low tumors (Figure 1C, Supplementary Figure 3). When pRPA2 foci were evaluated in the context of HR status, patients with RAD51-Low, pRPA2-High tumors had similar PFS (20.1 vs 14.4 months) as patients with RAD51-Low, pRPA2-Low tumors (Figure 1D). However, patients with RAD51-High, pRPA2-High tumors had longer PFS than patients with RAD51-High, pRPA2-Low tumors (15.2 vs 7.8 months) (Figure 1D).

**Figure 1.**
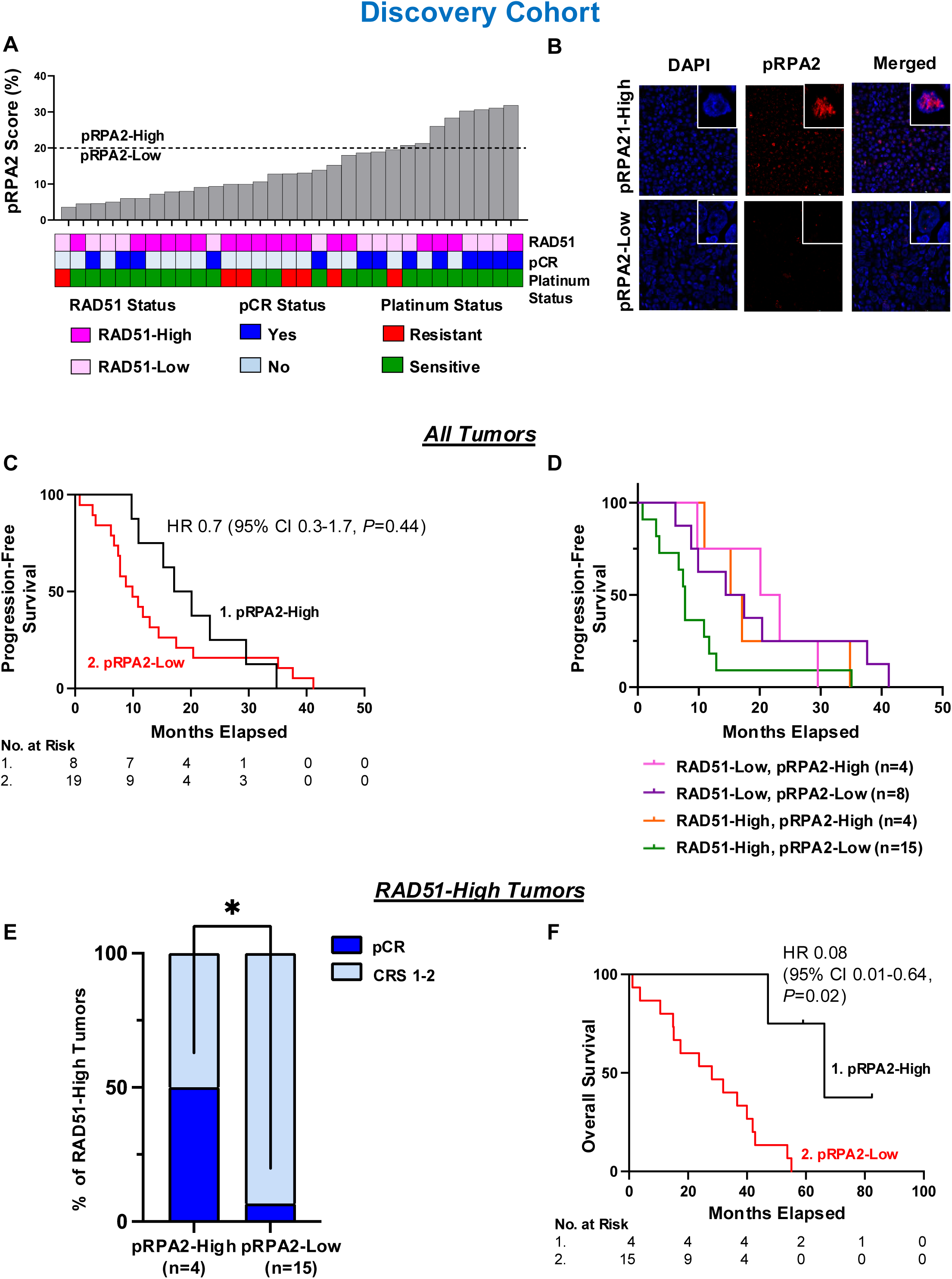
RAD51 and pRPA2 scores predict platinum chemotherapy response and survival in ovarian cancer in a discovery cohort. **A,** pRPA2 score, RAD51 score, pathologic complete response (pCR), and platinum sensitivity in patients with high-grade serous ovarian cancer before neoadjuvant chemotherapy. pRPA2 scores are defined as the percent of cells having ≥ 2 pRPA2 foci. Dotted black line indicates manual quantification 20% cut-off, which delineates pRPA2-High and pRPA2-Low tumors. All foci were counted in n>100 cells per experiment. Technical replicates were performed for 30% of samples. **B,** Representative images of DAPI and pRPA2 and overlay of DAPI/pRPA2 in patient-derived FFPE HGSOC sample. **C,** Kaplan Meier curves evaluating progression-free survival stratified by pRPA2 score in discovery cohort (n=27, HR 0.7, 95% CI 0.3-1.7, *P*=0.4). **D,** Progression-free survival in four RAD51/pRPA2 subgroups. **E,** Proportion of RAD51-High tumors that had a complete pathologic response (pCR) stratified by pRPA2 score *(P=*0.03) **F,** Kaplan Meier curves evaluating overall survival in RAD51-High tumors stratified by pRPA2 scores (n=19, HR 0.08, 95% CI 0.01-0.64, *P*=0.02*). *P*<0.05, by student’s two-tailed t-test.

To determine the ability of pRPA2 foci to directly predict platinum chemotherapy sensitivity in HR-proficient tumors, we assessed the association between pRPA2 foci and chemotherapy response score in RAD51-High tumors. A chemotherapy response score is assigned to each tumor at the time of interval cytoreductive surgery according to a validated histopathologic system and is a direct readout of platinum chemotherapy sensitivity (34). Thirteen (41.9%) tumors had a pathologic complete response (pCR): 10 (83.3%) RAD51-Low and 3 (15.8%) RAD51-High. Amongst the RAD51-Low tumors, there were no differences in pCR between pRPA2-High and pRPA2-Low tumors (Supplementary Figure 4A). Conversely, amongst RAD51-High tumors, pRPA2-High tumors had significantly higher rates of pCR than pRPA2-Low tumors (50% vs. 6%, *P*=0.03) (Figure 1E). In this group, pRPA2 foci predicted pCR to platinum chemotherapy with a specificity of 87.5% and a negative predictive value of 93.3%. Further, patients with RAD51-High, pRPA2-High tumors had longer OS than those with pRPA2-Low tumors (66.3 vs 28.1 months, *P*=0.02) (Figure 1F).

Because RAD51-High, pRPA2-High tumors exhibited similar platinum sensitivity as RAD51-Low tumors, we grouped RAD51-Low and RAD51-High, pRPA2-High tumors for further analysis. Patients with RAD51-Low tumors and those with RAD51-High, pRPA2-High tumors (n=16) were significantly more likely to have a pCR than those with RAD51-High, pRPA2-Low tumors (n=15) (74% vs. 6.7%, Relative Risk 11.3, 95% CI 1.7-76.3, *P*<0.001) (Supplementary Figure 4B). The combined assay predicted pCR to chemotherapy with 92.3% sensitivity, 77.8% specificity, 93.3% negative predictive value, and 75.0% positive predictive value. When controlled for stage and *BRCA1/2* mutation status, patients with RAD51-Low or RAD51-High, pRPA2-High tumors had significantly longer PFS (17.1 vs 7.7 months, *P*=0.02) (Supplementary Figure 4C) and OS (59.8 vs 28.1 months, *P*<0.001; Supplementary Figure 4D) than patients with RAD51-High, pRPA2-Low tumors.

To facilitate translation of the RAD51, pRPA2 assay into clinical care, we automated analysis of pRPA2 foci (automated analysis of RAD51 assay was described in (9)). To do so, we used linear coefficient of linear regression modeling to establish an automated pRPA2 score cutoff that corresponded to the manual cutoff of 20%. We found strong correlation between a manual cutoff of 20% and an automated pRPA2 cutoff of 16% (*r* = 0.7, *P <* 0.001; Cohen’s Kappa coefficient 0.72, *P* < 0.001). Using this cutoff, automated quantification had accuracy of predicting manual quantification of 96.6%, sensitivity of 100%, and specificity of 89.3%. Supplementary Figure 5 and Figure 2 illustrate the automated workflow and scoring criteria for our proposed combined HR and replication stress assay.

**Figure 2.**
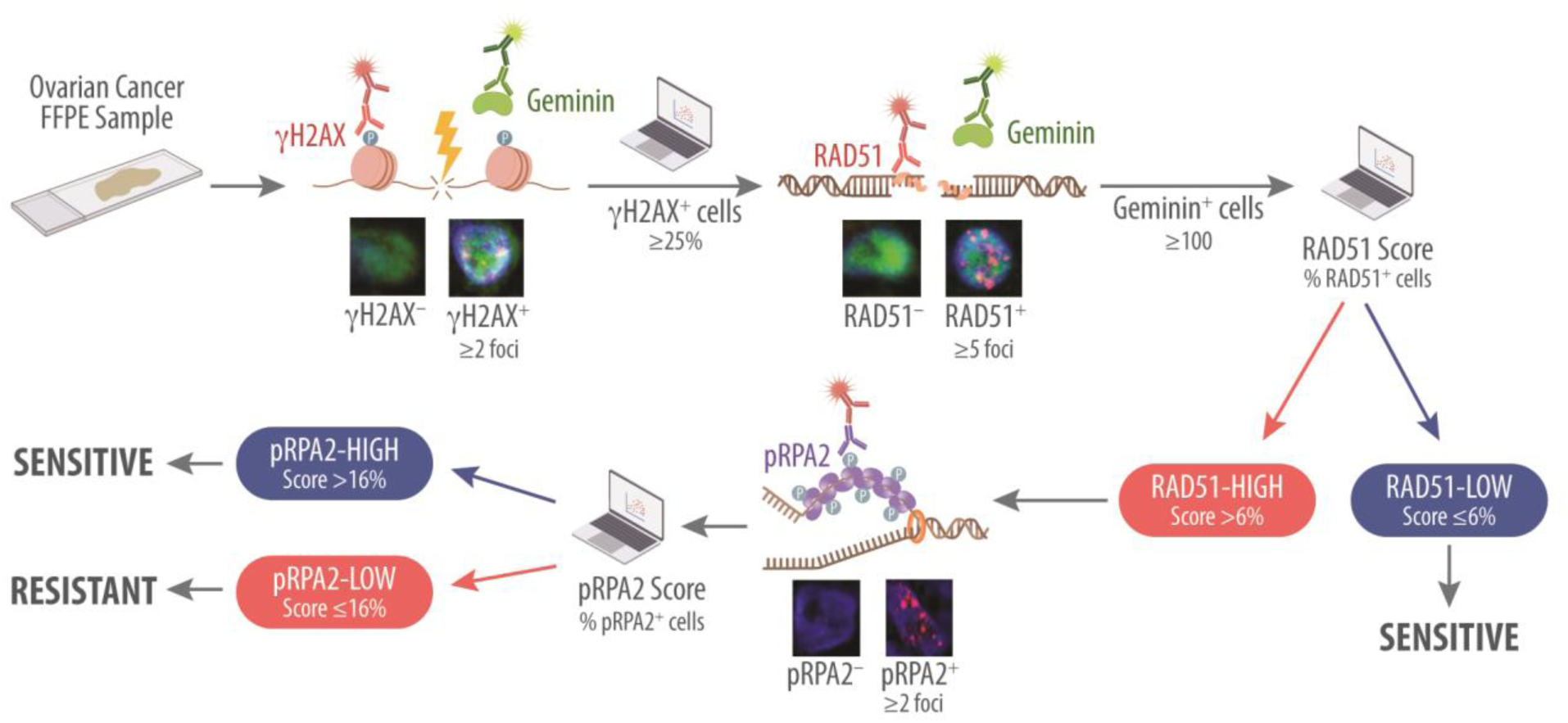
A functional assay to predict patient response to chemotherapy. Schematic of combined RAD51 and pRPA2 immunofluorescence assay in HGSOC FFPE samples including automated quantification. After confirming cellularity in FFPE ovarian cancer patient samples, the samples were screened for γH2AX. Only samples with 25% or more cells displaying two or more γH2AX foci were included in the analysis, ensuring sufficient DNA damage to elicit a response. RAD51 foci were then assessed in geminin-positive cells, with cells containing five or more RAD51 foci classified as positive. Samples were considered RAD51-High if 6% or more of the geminin-positive cells were positive for RAD51 on automated analysis. A minimum of 100 cells were assessed per sample. Subsequently, samples were evaluated for pRPA2 foci. Cells with two or more pRPA2 foci were considered positive, and samples with >16% of positive cells on automated analysis were considered positive.

To validate the automated combined RAD51 and pRPA2 foci assay to predict platinum responses, we screened an additional 246 high-grade serous ovarian, fallopian tube, or primary peritoneal tumors from patients receiving front-line platinum chemotherapy (Supplementary Figure 6). Two (0.8%) tumors were excluded because γH2AX scores were <25%, so 244 tumors were included in the final analysis (Table 1). All of these were high-grade serous ovarian cancer, and the majority were advanced stage disease (94.6%) and platinum sensitive (86.9%). One hundred twenty-two (50.5%) were pRPA2-High, and 120 (49.5%) were pRPA2-Low (Figure 3A). Of the 184 (75.4%) RAD51-High tumors, 96 (52.2%) were pRPA2-High, and 88 (47.8%) were pRPA2-Low.

**Figure 3.**
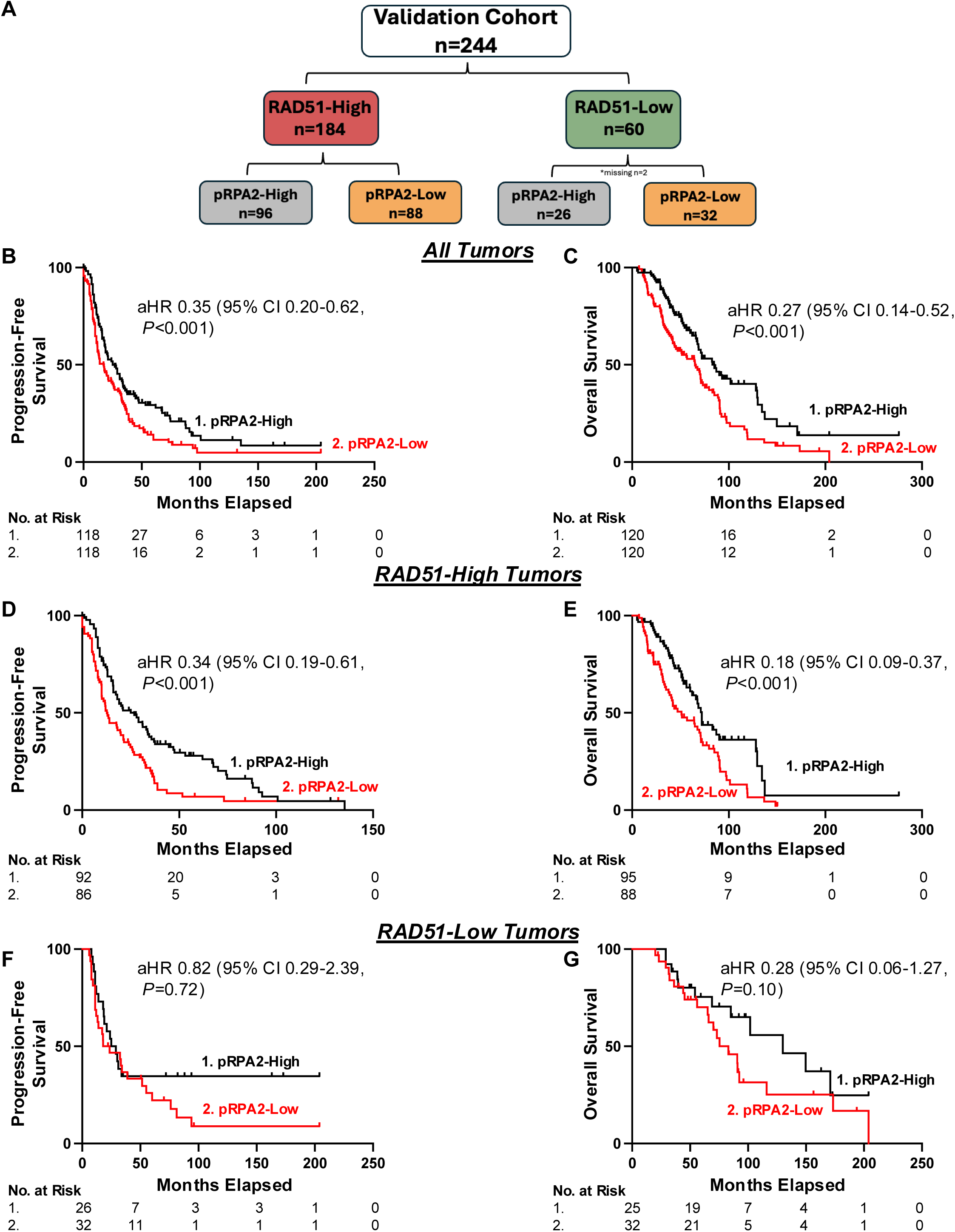
Automated pRPA2 and RAD51 scores predict platinum chemotherapy response and survival in a validation cohort. **A,** RAD51 scores and pRPA2 scores in the validation cohort. **B,** Kaplan Meier curves evaluating progression-free survival stratified by pRPA2 scores in validation cohort (n=236 aHR 0.35, 95% CI 0.20-0.62, *P<*0.001). ***C*,** Kaplan Meier curves evaluating overall survival stratified by pRPA2 score in validation cohort (n=240, aHR 0.27 95% CI 0.14-0.52, *P<*0.001). **D,** Kaplan Meier curves evaluating progression-free survival in RAD51-High tumors stratified by pRPA2 score (n=178, aHR 0.34, 95% CI 0.19-0.61, *P*<0.001). **E,** Kaplan Meier curves evaluating overall survival in RAD51-High tumors stratified by pRPA2 score (n=183, aHR 0.18, 95% CI 0.09-0.37, *P<*0.001). **F,** Kaplan Meier curves evaluating progression-free survival in RAD51-Low tumors stratified by pRPA2 score (n=58, aHR 0.82, 95% CI 0.29-2.39 *P*=0.72) **G,** Kaplan Meier curves evaluating overall survival in RAD51-Low tumors stratified by pRPA2 score (n=57, aHR 0.28, 95% CI 0.06-1.27, *P*=0.10). *aHR= adjusted hazard ratio for: age, stage, residual disease and *BRCA* status.

On multivariate analysis controlling for stage, age, residual disease at the time of cytoreductive surgery, and *BRCA1/2* mutation status, pRPA2 foci status was predictive of survival in the total cohort (Figure 3B-C). Upon further analysis, this effect seemed to be driven by the RAD51-High cases. Within the RAD51-High cases, pRPA2-High tumors had significantly longer PFS (26.9 vs. 12.7 months, *P*<0.001) (Figure 3D) and OS (72.0 vs. 51.0 months, *P*<0.001) (Figure 3E) than patients with pRPA2-Low tumors. Conversely, consistent with our findings in the discovery cohort, pRPA2 foci were not predictive of PFS or OS in RAD51-Low tumors (Figure 3F-G). These data suggested that replication stress is a major determinant of therapy sensitivity in HR-proficient tumors and can be implemented as a biomarker predictive of platinum chemotherapy response.

To confirm the clinical applicability of the combined biomarker, we evaluated survival after grouping RAD51-Low and RAD51-High, pRPA2-High tumors. One hundred fifty-five (63.5%) were RAD51-Low or RAD51-High, pRPA2-High, and 89 (36.5%) were RAD51-High, pRPA2-Low. RAD51-Low and RAD51-High, pRPA2-High tumors were more likely to have sustained responses to platinum chemotherapy (PFS ≥ 12 months) than RAD51-High, pRPA2-Low tumors (71.1% vs. 44.8%, Relative Risk 1.59, 95% CI 1.23-2.04, *P*<0.001). On multivariate analysis controlling for stage, age, residual disease at the time of cytoreductive surgery, and *BRCA1/2* mutation status, patients with RAD51-Low or RAD51-High, pRPA2-High tumors had significantly longer PFS (29 vs. 12 months, *P*<0.001) (Supplementary Figure 7A) and OS (83 vs. 47 months, *P*<0.001) (Supplementary Figure 7B) than patients with RAD51-High, pRPA2-Low tumors.

### RAD51 and pRPA2 foci predict survival outcomes in RAD51-High tumors after PARP inhibitor treatment

We next wanted to determine the utility of pRPA2 foci for predicting survival after PARP inhibitor treatment in RAD51-High tumors. We analyzed samples from a cohort of 87 ovarian cancer patients who received PARP inhibitors at any point during their treatment course (Table 1, Figure 4A-B). These patients were from two independent cohorts with median follow-up times of 51.7 ± 24.2 months (n=60) and 74.7 ± 36.7 months (n=26). Sixty-six (75.9%) tumors were RAD51-High. Of these, 41 (62.1%) were pRPA2-High and 25 (37.9%) pRPA2-Low. Thirty-five (40.2%) of patients received PARP inhibitors as upfront maintenance, 28 (32.3%) as recurrent therapy, and 24 (27.6%) the timing of therapy was unknown (Figure 4B). In the PFS analysis, we only included the 25 patients with RAD51-High tumors who received PARP inhibitors as upfront maintenance. In these 25 patients, those with pRPA2-Low tumors had a median PFS of 35.1 months whereas those with pRPA2-High tumors did not meet a median survival (64.3% of cases were censored because they had not yet recurred) (Figure 4C). This difference was clinically profound, but not statistically significant as the data are not yet mature. We next examined OS in the 66 patients with RAD51-High tumors who received a PARP inhibitor at any time. On multivariate analysis controlling for stage, age, residual disease at the time of cytoreductive surgery, and *BRCA1/2* mutation status, patients with RAD51-Low or RAD51-High, pRPA2-High tumors had significantly longer OS than those with RAD51-High, pRPA2-Low tumors (87.0 vs. 42.1 months, *P*<0.001) (Figure 4D).

**Figure 4.**
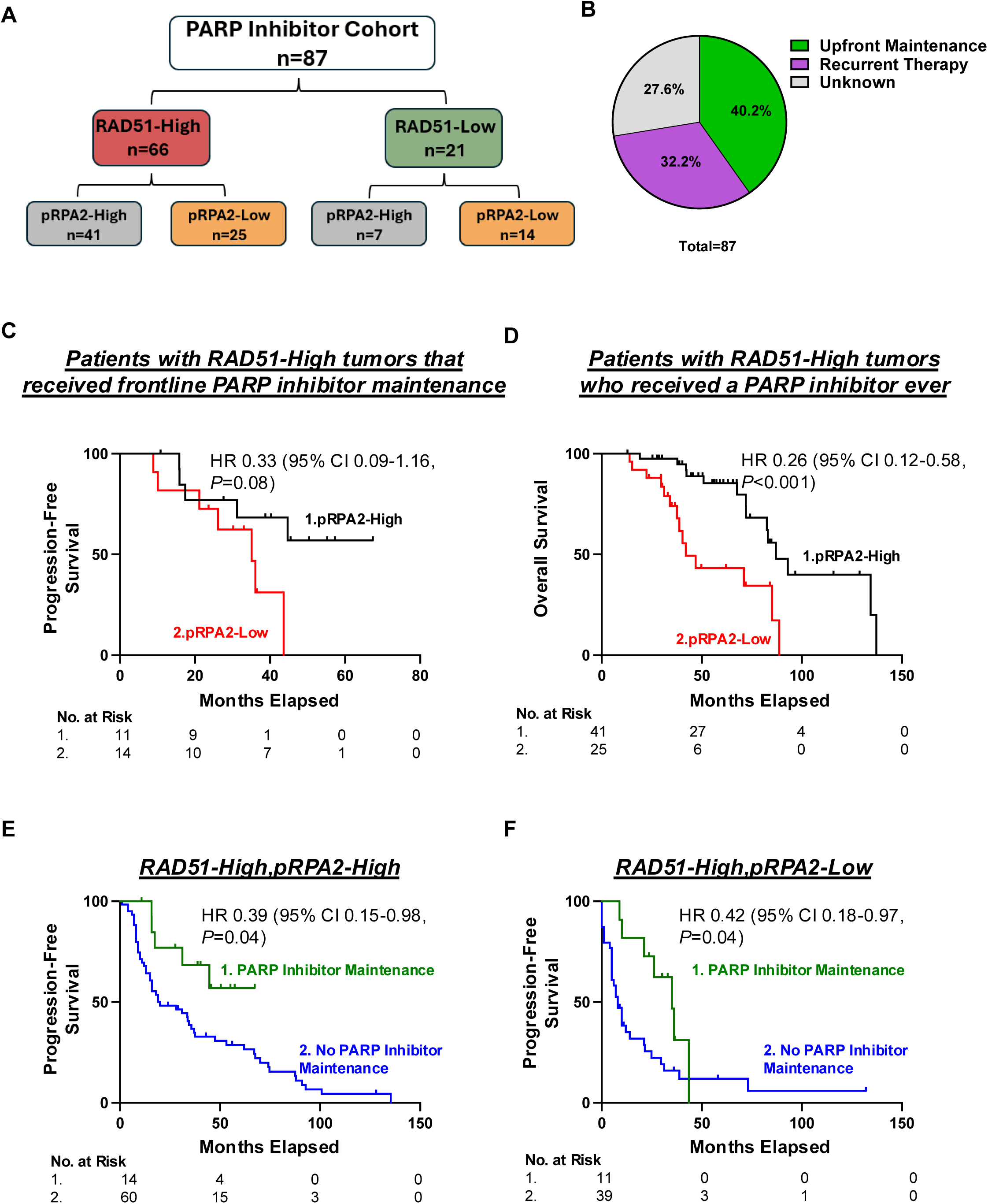
Automated pRPA2 and RAD51 scores predict survival in patients with high grade serous ovarian cancer (HGSOC) receiving PARP inhibitors. **A,** RAD51 scores and pRPA2 scores in the validation cohort of patients who received a PARP inhibitor at any time during their treatment (n=87). **B,** Pie chart illustrating the timing of PARP inhibitor therapy during a patient’s treatment course **C,** Kaplan Meier curves evaluating progression-free survival in with RAD51-High HGSOC who received a frontline PARP inhibitor stratified by pRPA2 scores (n=25, HR 0.33, 95% CI 0.09-1.16, *P=*0.08). **D,** Kaplan Meier curves evaluating overall survival in tumors with RAD51-High tumors who received PARPi at any point during their treatment stratified by pRPA2 scores (n=66, HR 0.26, 95% CI 0.12-0.58, *P<*0.001). **E,** Kaplan Meier curves evaluating progression-free survival in tumors with RAD51-High, pRPA2-High scores stratified by whether or not they received frontline PARP inhibitor maintenance therapy (n=74, HR 0.39, 95% CI 0.15-0.98, *P*=0.04). **F,** Kaplan Meier curves evaluating progression-free survival in tumors with RAD51-High, pRPA2-Low scores stratified by whether they received frontline PARP inhibitor maintenance therapy (n=50, HR 0.42, 95% CI 0.18-0.97, *P*=0.04).

Next, we analyzed PFS in patients with RAD51-High tumors who did and did not receive upfront PARP inhibitor maintenance stratified by pRPA2 status. We evaluated patients with RAD51-High, pRPA2-High tumors. At the time of analysis (median follow-up 54 months), only 35.7% of patients with pRPA2-High tumors who received PARP inhibitor maintenance had a recurrence. Conversely, 86.7% of those who received no PARP inhibitor maintenance group had a recurrence. PARP inhibitor maintenance in these patients was associated with longer PFS (undefined vs 20.0 months, *P*=0.04) (Figure 4E) and a trend towards OS (82.5 vs 67.6 months, *P=*0.09) (Supplemental Figure 8A). Notably, in the OS analysis, 92.9% of patients who received PARP inhibitor maintenance were censored. Next, we analyzed RAD51-High, pRPA2-Low tumors. In these patients, PARP inhibitor maintenance was associated with increased PFS (35.1 vs 8.0 months *P=*0.04) (Figure 4F), but not with OS (37.6 vs 33.0 months, *P*=0.26) (Supplemental Figure 8B). Markedly, the median PFS in patients with RAD51-High, pRPA2-High tumors treated with PARP inhibitors was significantly longer than in patients with pRPA2-Low tumors. In total, this data suggests that PARP inhibitor maintenance significantly benefits patients with RAD51-High, pRPA2-High tumors.

### RAD51 and pRPA2 foci predict patient survival in recurrent tumors

A major limitation of currently available genomic HR assays is their static nature. The genomic instability that results from HR-deficient DNA repair mechanisms is not reversible, so a tumor will be defined by genomic scar assays as HR-deficient even if it evolves to be HR-proficient (35-37). In our assay, RAD51 and pRPA2 staining only produces foci if the tumor cells recently experienced HR and replication stress thereby providing a dynamic reading of HR. To test this idea, we evaluated tumor RAD51 and pRPA2 scores in 42 patients from whom we had recurrent tumor biopsies. Of these, tumors from 4 (9.5%) patients had γH2AX scores < 25% and were excluded from the analysis. Of the remaining 38 patients, 37 had matched tumor samples from diagnosis and recurrence (Figure 5A, Supplemental Figure 6). At the time of diagnosis, 29 (76.3%) tumors were RAD51-High and 8 were RAD51-Low (21.1%) (Figure 5B). At the time of recurrence, 14 (37.8%) tumors changed RAD51 classification. Patients with RAD51-Low tumors at recurrence had longer survival after recurrence than those with RAD51-High tumors (53.5 vs. 23.9 months, *P*=0.06) (Figure 5C). Next, we considered pRPA2 score. Ten (27.0%) patients had a change in their pRPA2 classification between diagnosis and recurrence (Figure 5D). At the time of recurrence, 9 (24.3%) were pRPA2-High and 28 (75.7%) were pRPA2-Low (Figure 5D). To improve the power of this analysis, we next considered RAD51-Low and RAD51-High, pRPA2-High tumors together. Twenty (54.0%) tumors had the same classification at diagnosis and recurrence, and 17 (45.9%) were reclassified (Figure 5B, E). Patients with RAD51-Low or RAD51-High, pRPA2-High tumors at recurrence had longer OS (50.9 vs. 23.2 months, *P=*0.01) than those with RAD51-High, pRPA2-Low tumors (Figure 5F). We conclude that the RAD51, pRPA2 assay is dynamic and useful at time of recurrence.

**Figure 5.**
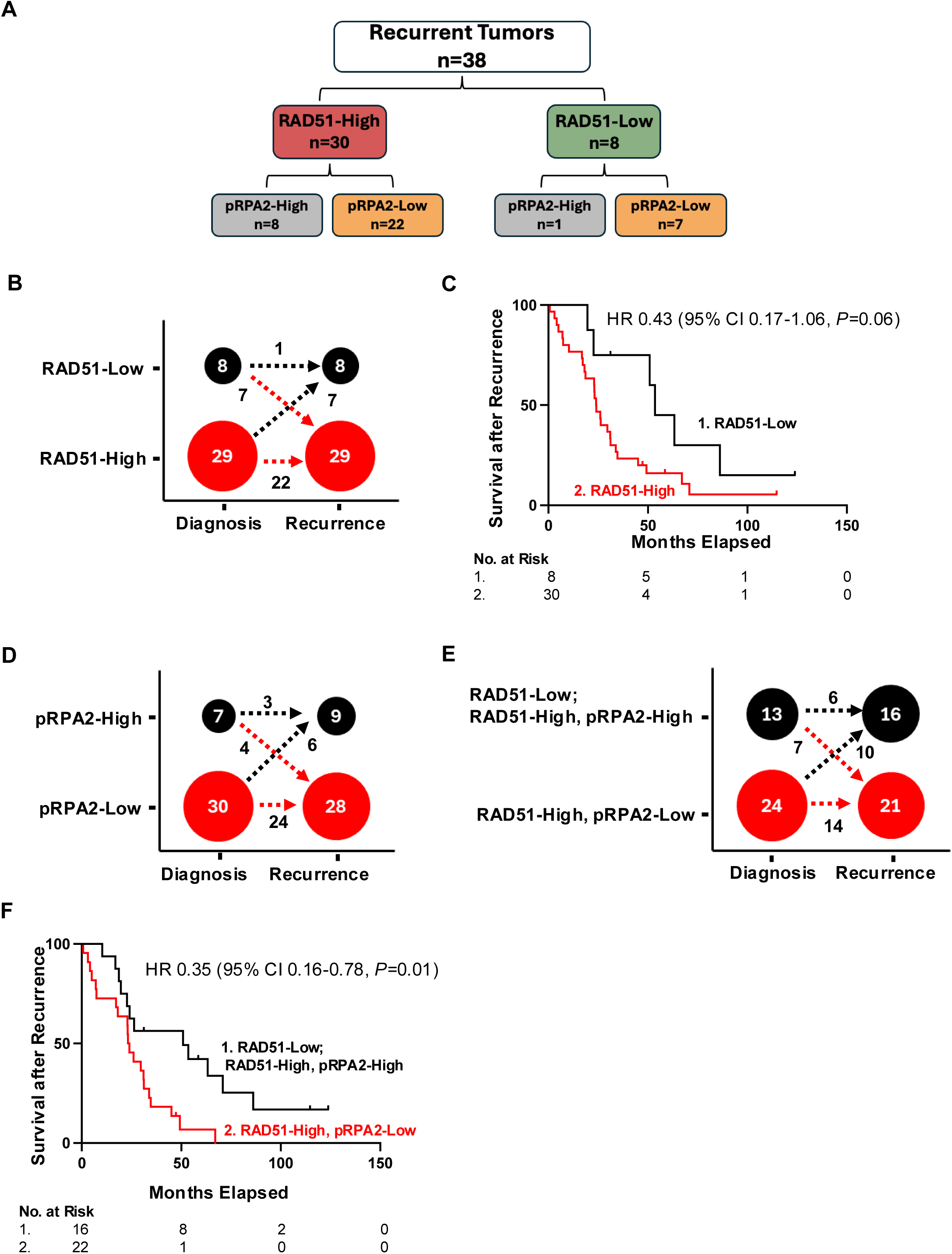
Automated pRPA2 and RAD51 at the time of recurrence predict survival. **A,** RAD51 scores and pRPA2 scores in recurrent tumors (n=38). **B,** Stratifying tumors by RAD51 score at time of diagnosis and score at time of recurrence (n=37). **C,** Kaplan Meier curves evaluating overall survival of recurrent patient samples by RAD51 scores (n=38, HR 0.43, 95% CI 0.17-1.07, *P=*0.07). **D** Stratifying tumors by pRPA2 score at time of diagnosis and score at time of recurrence (n=37). **E,** Stratifying patients by RAD51/pRPA2 score at time of diagnosis and score at time of recurrence (n=37). **F**, Kaplan Meier curves evaluating overall survival of recurrent patient samples by RAD51/pRPA2 scores (n=38, HR 0.35, 95% CI 0.16-0.78, *P=*0.01).

## DISCUSSION

While nearly all HR-deficient tumors are sensitive to therapy, a subset of HR-proficient tumors also exhibit therapeutic sensitivity (9). Here, we demonstrate that a key determinant of response in HR-proficient tumors is replication stress. To date, no functional biomarkers of replication stress have been developed for use on clinically available specimens. Our work describes the development and validation of a functional assay to measure replication stress in FFPE ovarian tumors. We first demonstrate that we can reliably evaluate replication stress in FFPE tumor cells with an antibody targeting Thr21 phosphorylated RPA2 (pRPA2). Next, in a discovery cohort, we show that RAD51-High (HR-proficient) tumors with high pRPA2 respond to platinum chemotherapy similarly to RAD51-Low (HR-deficient) tumors. We then used a validation cohort to demonstrate that patients with RAD51-High, pRPA2-High tumors had longer progression-free and overall survival after platinum chemotherapy than patients with RAD51-High, pRPA2-Low tumors. Additionally, our data suggest that pRPA2 foci can accurately identify which RAD51-High tumors will and will not respond to PARP inhibitors. Finally, we showed that the assay is dynamic and predictive in recurrent tumors.

Several lines of published and preliminary evidence indicate that increasing replication stress can sensitize HR-proficient tumors to DNA damaging therapy. First, replication stress-inducing agents, such as inhibitors of ATR, WEE1, and CHK1/2, are being explored in clinical trials with promising early results specifically in HR-proficient cancers (38-41). Second, in a phase II clinical trial in platinum-resistant ovarian cancer evaluating the combination of the replication stress-inducing agents gemcitabine and an ATR inhibitor, tumors with high replication stress responded better to combination therapy independent of HR status (38). Third, multiple studies have evaluated the combination of gemcitabine and carboplatin in platinum-sensitive and platinum-resistant ovarian cancer. They have demonstrated impressive response rates of over 75% in platinum-sensitive tumors and 65% in platinum-resistant tumors, with survival greater than 25 months in select platinum-resistant patients (42-47). This is remarkable for a cohort of patients who typically have response rates to subsequent chemotherapy of 15% and survival of 12 months.

Although our data illustrate a clear association between therapy response and pRPA2 foci, the mechanisms driving high pRPA2 in some tumors are not fully elucidated. High numbers of pRPA2 foci could reflect increased replication fork stalling, fork degradation, or ssDNA gaps. Fork stalling can result from DNA damage or replication obstacles. Upon encountering a DNA lesion, the replication polymerase decouples from the helicase, promoting DNA unwinding and accumulation of single-stranded DNA (ssDNA) between the polymerase and the helicase. This ssDNA is coated by RPA (25), which protects the ssDNA from nucleolytic degradation. Cells that can resolve this stalling and restart replication are often therapy resistant (48). Upon stalling, the replication fork can reverse. In this configuration, the complementary leading and lagging nascent strands of DNA anneal, creating a "chicken foot" structure (27). This mechanism relies on the ability of cells to protect the newly synthesized DNA by loading RAD51. If the cell cannot protect the DNA, nucleases degrade the DNA, leaving behind ssDNA that is coated with RPA. Replication fork degradation is associated with sensitivity to chemotherapy in established ovarian cancer cell lines (28). Alternatively, to overcome fork stalling, the replisome can skip the obstructing lesion, reprime the DNA with PRIMPOL, and continue replication. This results in a ssDNA gap, which is coated with RPA. Increased gaps can also be a result of defective Okazaki fragment processing or defective gap repair by translesion synthesis or template switching. Replication ssDNA gaps have been proposed to drive therapy sensitivity independent of HR status in cell lines (23, 24, 29). It is possible that increased pRPA accumulation can be a cumulative event from varied mechanisms of replication fork remodeling. Further work is necessary to determine which events are responsible for therapy response in HR-proficient tumors.

An important finding of our study is that the RAD51, pRPA2 assay might be useful for recurrent tumor samples. HR status is currently assessed clinically by germline or somatic mutation testing or evaluation of genomic scars. However, these assays all provide a snapshot of past events and do not necessarily reflect the current HR capacity of a tumor. Clinical ability to predict whether tumors will respond to platinum at recurrence is also poor. For example, historical data suggest that cancers in up to 50% of patients in a “platinum-resistant” cohort respond to further platinum treatment (49). Conversely, cancers in only about 50-60% of patients with disease-free intervals greater than 6 months – considered “platinum-sensitive” – respond to repeat platinum-based therapy (19). Therefore, over 50% of patients have misclassified platinum sensitivity and are either not offered the most effective drug for their disease or are given an ineffective, toxic therapy (49). With additional validation, the RAD51, pRPA2 assay described here may allow clinicians to expand chemotherapy to patients with recurrent disease and improve their long-term outcomes.

Our work has several limitations. First, although automated, our assay still requires a trained technician to image the stained microscopy slides. Second, our assay uses deconvolution imaging, which might not be readily available. Third, we did not have genomic scar data for these tumors. Thus, we were unable to directly compare the performance of the RAD51, pRPA2 assay to that of genomic scars. This is a future aim of our work. Fourth, the survival data for patients who received PARP inhibitors was not mature, so the differences described are likely underestimated. Fifth, we observed considerable tumor heterogeneity in RAD51 and pRPA2 foci staining within tumor samples. While our results suggest that this heterogeneity can be addressed by including over 100 cells in each analysis, varying degrees of heterogeneity might carry clinical implications that were beyond the scope of this study to assess. And finally, while this assay predicts therapy resistance, it does not propose alternative treatment regimens. However, this work does establish a clear mechanism of sensitivity in HR-proficient tumors which can be targeted to overcome resistance. For example, perhaps patients with pRPA2-Low tumors should be treated with platinum chemotherapy in combination with a replication stress inducing agent such as gemcitabine. On the other hand, those with pRPA2-High tumors will respond well to standard of care platinum chemotherapy and paclitaxel. Future work will clarify this.

In conclusion, our combined, automated RAD51 and pRPA2 assay can be a powerful clinical tool for predicting response to platinum chemotherapy and PARP inhibitors. Its dynamic nature allows for real-time tumor behavior evaluation, making it particularly valuable in managing recurrent disease. Further prospective studies are warranted to validate these findings and integrate the assay into routine clinical practice.

## METHODS

### Sex as biological variable

Our study exclusively examined females because ovarian cancer is only relevant in females.

### Patient Samples/Study Approval

For the discovery cohort, tissue was collected prospectively from patients with Stage III-IV high-grade serous ovarian cancer before neoadjuvant chemotherapy as part of a national clinical trial (IRB #201407156). All tissues were collected between December 2014 and December 2018. Chemotherapy response score was assigned at the time of interval cytoreductive surgery according to a validated system (50). A chemotherapy response score of 3 was consistent with a pathologic complete response (pCR), and a score of 1-2 was considered a poor response to platinum chemotherapy. Progression-free survival (PFS) and overall survival (OS) were calculated from the time of interval cytoreductive surgery.

For the validation cohort, four tissue microarrays of high-grade serous ovarian cancer tumor samples were used. The first was comprised of deidentified samples from the University of Kansas. The second and third were built in Anatomic and Molecular Pathology Core Laboratories at Washington University (IRB# 202301067). These both contained primary and/or metastatic tumors collected during primary or interval cytoreductive surgery after chemotherapy. The fourth, from Cedars-Sinai Medical Center, contained deidentified primary, metastatic, and recurrent tumor samples (IRB: Pro44852) (51).

### Development of primary ovarian cancer cells (POVs) and cell culture/Study Approval

Tissues were prospectively collected for our Gynecologic Oncology biorepository (IRB #201105400 and #201706151) with informed patient consent. To establish primary cell lines, ascites was collected from patients with advanced-stage high-grade serous ovarian cancer (HGSOC) and transferred to culture flasks containing 1:1 (V/V) RPMI supplemented with 20% FBS and 1% penicillin and streptomycin. After 1 to 2 weeks, attached and proliferating cells were passaged and used for experiments. Cells were discarded after 1 to 2 passages.

### Immunofluorescence

POVs (40,000 cells per well) were plated in 8-well chamber slides. Cells were washed with cold PBS, fixed for 10 min with 2% paraformaldehyde, permeabilized for 20 min with 0.2% Triton X-100 in PBS, and washed again with PBS. Cells were blocked for 30 minutes in PBS, 0.5% BSA, 0.15% glycine, and 0.1% Triton X-100 and incubated overnight at 4 °C with primary antibodies (Supplementary Table 1) in staining buffer. Cells were then stained with secondary antibodies (Supplementary Table 1), followed by DAPI (Sigma). Cells were imaged on a Leica TCS SPE inverted confocal microscope. Raw images were exported, and JCountPro was used to count the foci (9, 52, 53). At least 100 cells were analyzed for each treatment group in duplicate.

### Transfection

TYKNU and OVCAR8 cells were transfected with RPA2 Silencer Select siRNA (Thermo Fisher Scientific) or non-targeting control siRNA (Thermo Fisher Scientific) using Lipofectamine RNAiMAX transfection reagent (Thermo Fisher Scientific) according to the manufacturer’s instructions. Two days after transfection, cells were treated with hydroxyurea (Sigma).

### FFPE cell blocks

The cells were fixed in 4% paraformaldehyde for 1 hour. The cell pellet was embedded in 2% agarose and put into a cassette in 10% formalin for an additional 24 hours. The cell pellets were washed with deionized water and dehydrated in sequential concentrations of ethanol (30%, 50%, and 70%). The samples were embedded in paraffin wax, cut into 4-mm sections, and mounted onto slides for further analysis.

### Western blot analysis

Cells transfected with siRPA2 or non-targeting siRNA (siControl) were lysed in 9 mol/L urea, 0.075 mol/L Tris, pH 7.6, and protein concentration was determined by using the Bradford assay (Bio-Rad). Protein lysates (60– 100 µg) were separated on SDS-PAGE, transferred to nitrocellulose, probed with primary antibody (Supplementary Table 1) at 4 °C overnight in humid chambers, washed, and incubated with corresponding horseradish peroxidase–conjugated secondary antibodies (Jackson ImmunoResearch). Pierce ECL Western Blotting Substrate (Thermo Scientific) was used to detect signal, and chemiluminescence was measured on a ChemiDoc (Bio-Rad Laboratories).

### MTS cell viability assay

Cells were seeded at 2000 cells per well in a 96-well plate. After 24 hours, cells were treated with a range of carboplatin (Teva pharmaceuticals). Viability was assessed 6 days post-treatment by using an MTS/PMS (Promega #PR-G1112) solution and reading absorbance at 490 nm on an Infinite M200 Pro plate reader (Tecan, Inc.). IC_50_ values were calculated in GraphPad Prism (GraphPad Software, Inc.).

### FFPE immunofluorescence

Immunofluorescence on FFPE samples was completed as previously described (9). Briefly, hematoxylin and eosin-stained slides were examined by a board-certified pathologist to assess cellularity and diagnosis. Corresponding unstained slides (4 µm thick sections) were deparaffinized, dehydrated, and stained with primary and secondary fluorescent antibodies (Supplementary Table 1). Samples were stored at -20 °C. Imaging was performed on a Leica DMi8 microscope with Thunder imaging computational clearing.

### Automated foci analysis

Stained microscopy images were imported into R environment. After denoising, smoothing, and thresholding, a 2-dimensional convolution was applied to segment all the foci in the image. Then, the foci-positive cells were counted, and the ratio of foci-positive cells to all cells was calculated. With multiple images from each patient (n ≥ 5), a median ratio was computed to estimate the γH2AX, RAD51, and pRPA2 scores. The linear coefficient of linear regression modeling was used to determine automated cutoffs.

### Statistical analysis

Traditional statistical analyses were performed in GraphPad Prism 9 and SPSS version 27 software (9). Baseline patient demographics were assessed with descriptive statistics, excluding any missing data from the analysis. Independent Student’s t-tests and Mann-Whitney U tests were used to compare continuous variables. One-way ANOVA was applied where appropriate. For correlation analyses, the Pearson product-moment correlation coefficient was used. Poisson regression was used to determine relative risks (RRs). PFS and OS were measured from date of surgery to date of evidence of disease recurrence, death, or last contact if no recurrence was observed. Patients without recurrence or still alive were censored at their last contact date. The Kaplan-Meier method estimated survival times, with comparisons made via the log-rank test. Cox proportional hazards regression was done in univariate and multivariate formats as needed. Statistical significance was set at P < 0.05, with two-tailed 95% confidence intervals.

### Sample size calculation

The sample size necessary to detect a significant Hazard Ratio (HR) of PFS from the patients with RAD51-Low or RAD51-High, pRPA2-High tumors compared to the patients with RAD51-High, pRPA2-Low tumors (54, 55) was calculated. If the expected HR of RAD51-Low or RAD51-High, pRPA2-High tumors was less than 0.3 based conservatively on our preliminary data, then 26 patients would provide the study with 80% power or 34 patients with 90% power, at a two-sided significance level of 0.05. Therefore, the validation cohort of 244 patients was highly adequate to assess the predictive value of the RAD51, pRPA2 assay.

## Data availability

The data produced in this study can be made available upon request. For inquiries, please contact the corresponding author.

## AUTHOR CONTRIBUTIONS

AS and AC conceptualized experiments, conducted experiments, acquired data, analyzed data, write, edit and review manuscript

RK, ML, KR, JB acquired data, edited the manuscript, and provided critical feedback and edits on manuscript SH provided patient samples and edited manuscript

CS acquired data and edited manuscript

BS, LK, CM, AH, PT, DM and MP provided patient samples and critical feedback and edits on manuscript VS provided experimental design advice and provided critical feedback and edits on manuscript

IH evaluated patient samples and provided critical feedback and edits on manuscript

AW, BK, SO, and KF provided patient samples and provided critical feedback and edits on manuscript LS analyzed and confirmed pathology of samples and provided critical feedback and edits on manuscript

PV conceptualized and designed experiments and provided critical feedback and edits on manuscript

EL conceptualized experiments, conducted experiments, acquired data, analyzed data, wrote manuscript and edited manuscript

PZ wrote code for automatic quantification of foci, conducted statistical analyzes and provided critical feedback and edits on manuscript

DK proved patient samples, experimental design feedback, and provided critical feedback and edits on manuscript

MM provided funding and reagents, conceptualized experiments, conducted experiments, acquired data, analyzed data, wrote and edited manuscript

## ACKNOWLEDGEMENTS

We especially wish to acknowledge the patients who generously donated their tumors specimens to make this work feasible. We’d also like to acknowledge Deborah Frank for her manuscript editing, Davi Martins for his manuscript editing, Thomas Walsh for contributing to the tissue microarrays, KIMEN Design 4 Research for generating illustrations, and Mr. Glen Huns for support of this project.

We also wish to recognize the following entities for their support of this project: Reproductive Scientist Development Program (RSDP) and the Gynecologic Oncology Group Foundation (MM), Pilot Translational and Clinical Studies function of the Washington University Institute of Clinical Translational Sciences and the Foundation for Barnes-Jewish Hospital CTRFP1719 (MM), The Cancer Biology Pathway Training Grant 5T32CA113275-17 (AS), The Lucy, Anarcha, and Betsey (L.A.B.) Award from the Department of Obstetrics and Gynecology at Washington University School of Medicine (AC), VA-ORD I01BX006020 (SO), and the National Institute of Health under award number R01CA243511 (DK). The content is solely the responsibility of the authors and does not necessarily represent the official views of the National Institutes of Health.

