## Supplemental Figures for "Replication stress marker phospho-RPA2 predicts response to platinum and PARP inhibitors in homologous recombination-proficient ovarian cancer"

**Supplementary Table 1. Summary of Antibodies**

| Antibody | Manufacturer/Catalog Number | Application | Dilution |
|---|---|---|---|
| RAD51 | Abcam/ab133534 | Immunofluorescence | 1:1000 |
| γH2AX | Millipore-Sigma/05-636 | Immunofluorescence | 1:500 |
| Phosphorylated RPA2 (Thr21) | ThermoFisher/BS-5693R | Immunofluorescence | 1:500 |
| Geminin | Leica, Novacastra/NCL-L Geminin | Immunofluorescence | 1:60 |
| Geminin | Proteintech/10802-1-AP | Immunofluorescence | 1:400 |
| Alexa Fluor 568, Anti-Ms | Invitrogen, A10037 | Immunofluorescence | 1:500 |
| Alexa Fluor 568, Anti-Rb | Invitrogen, A11011 | Immunofluorescence | 1:500 |
| Alexa Fluor 488, Anti-Rb | Invitrogen, A32731 | Immunofluorescence | 1:500 |
| Alexa Fluor 488, Anti-Ms | Invitrogen, A32723 | Immunofluorescence | 1:500 |
| Alexa Fluor 647, Ant-Rb | Invitrogen, A32733 | Immunofluorescence | 1:500 |
| Phosphorylated RPA2 (Thr21) | ThermoFisher/BS-5693R | Western blot | 1:1000 |
| HSP70 | ThermoFisher/PA5-34772 | Western Blot | 1:5000 |

**A**

| | Total (n=9) |
|---|---|
| Age (years) | 60.4±12.4 |
| FIGO Stage | |
| IIIA | 1 (11) |
| IIIB | 2 (22) |
| IIIC | 5 (56) |
| Unknown | 1 (11) |
| Histology | |
| Serous | 7 (78) |
| Endometrioid | 1 (11) |
| Clear Cell | 1 (11) |
| Unknown | 1 (11) |
| BRCA Mutation | |
| Yes | 1 (11) |
| No | 6 (67) |
| Unknown | 2 (22) |
| Prior Lines of Chemotherapy | |
| 0 | 6 (67) |
| 1 | 2 (22) |
| Unknown | 1 (11) |
| Platinum Response | |
| Sensitive (PFI≥6 months) | 3 (33) |
| Resistant (PFI≤6 months) | 4 (44) |
| Refractory (PFI≤4 weeks) | 1 (11) |
| Unknown | 1 (11) |

Data are n (%) unless stated otherwise. ± denotes standard deviation. PFI, Progression free interval

**B**

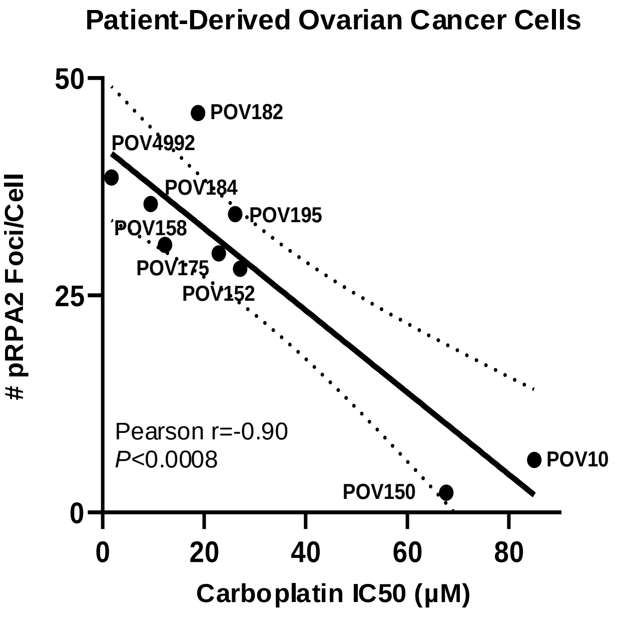

**Supplementary Figure 1. pRPA2 foci correlate with platinum chemotherapy response in patient-derived ovarian cancer cells**. **A,** Clinical demographics of patient-derived ovarian cancer cells. **B,** The IC50 of carboplatin in 9 POVs was determined and compared to baseline pRPA2 foci of the cells (Pearson r= -0.90, P<0.001).

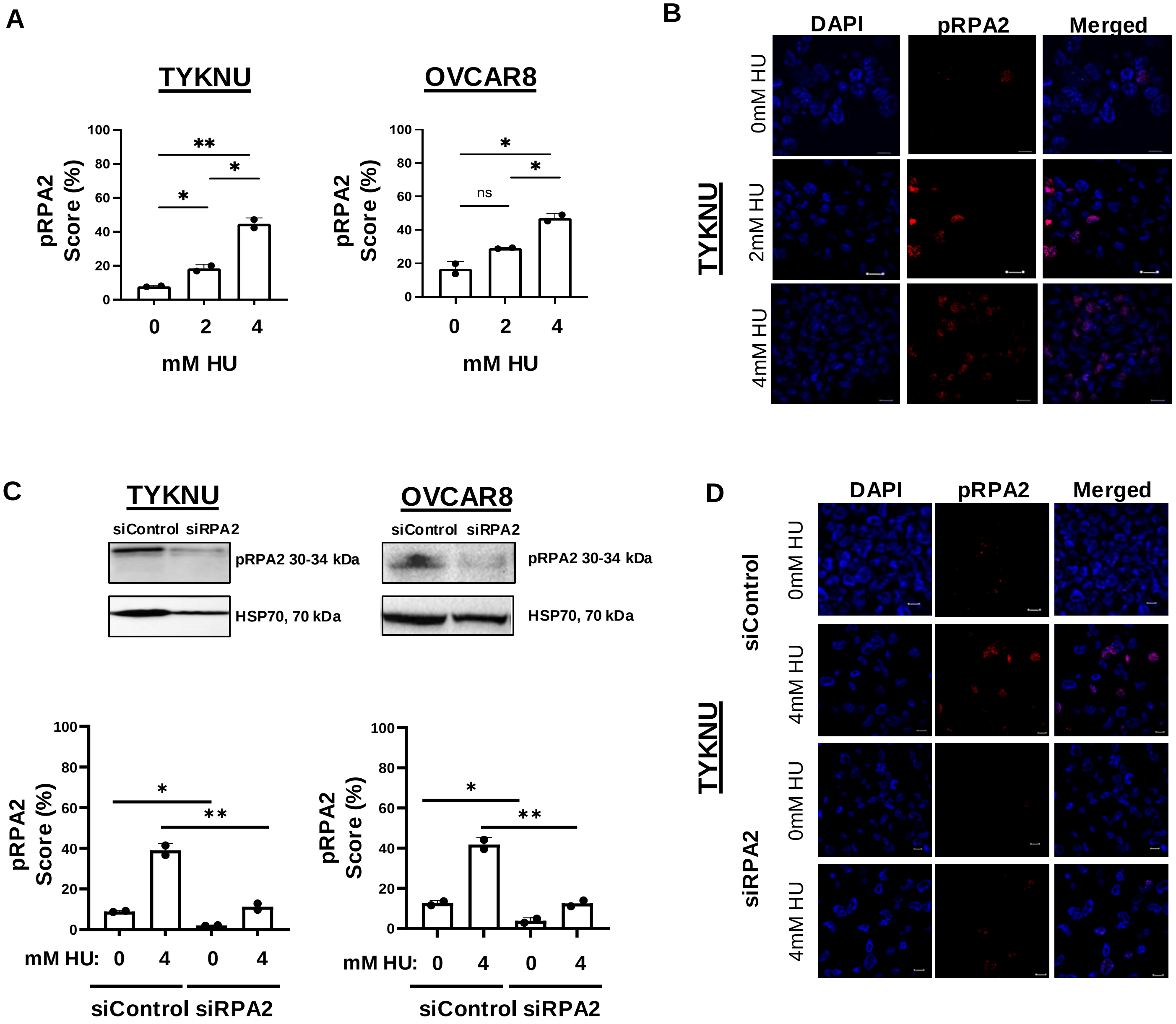
A
TYKNU
OVCAR8
pRPA2 Score (%)
mM HU
ns
B
DAPI
pRPA2
Merged
TYKNU
0mM HU
2mM HU
4mM HU
C
siControl
siRPA2
pRPA2 30-34 kDa
HSP70, 70 kDa
mM HU:
D
siControl
siRPA2

**Supplementary Figure 2. Validation of pRPA2 immunofluorescence assay in formalin-fixed paraffin-embedded (FFPE) samples. A,** Dynamic range of pRPA2 scores in two FFPE HR-proficient HGSOC cell lines after treatment with 0, 2, and 4 mM of hydroxyurea (HU) to induce replication stress. Cells were treated, fixed, embedded, and cut into 4 µm sections for evaluation. **B,** Representative images of DAPI, pRPA2 and colocalization of DAPI/pRPA2 at 63X in an FFPE HGSOC cell line treated with varying levels of hydroxyurea. Scale bars: 50 µm. **C.** Western Blot of HGSOC cell lines after transfection with siRNA targeting RPA2. pRPA2 scores in two HR-proficient HGSOC cell lines after transfection with short-interfering RNA (siRNA) targeting RPA2 (siRPA2) or a non-coding region (siControl) and exposed to 0 and 4 mM of hydroxyurea. Cells were treated, fixed, embedded, and cut into 4 µm sections for evaluation. **D.** Representative images of DAPI, pRPA2 and overlay of DAPI/pRPA2 at 63X in a FFPE HGSOC cell lines. **P*<0.05, ***P*<0.01, ****P*<0.001, *****P*<0.0001 by student's two-tailed t-test.

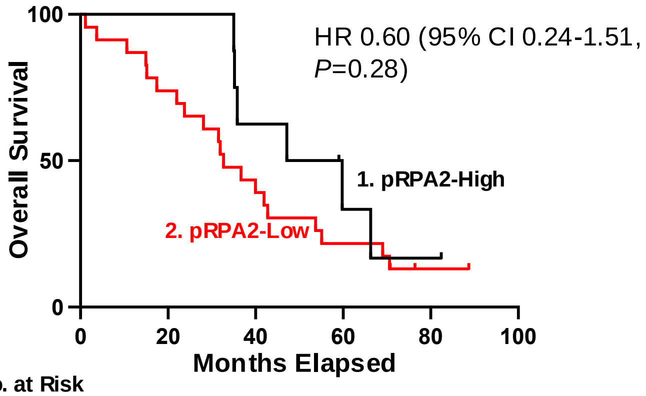


| **No. at Risk** | | | | | | |
|---|---|---|---|---|---|---|
| 1. | 8 | 8 | 5 | 2 | 1 | 0 |
| 2. | 23 | 17 | 9 | 5 | 1 | 0 |

**Supplementary Figure 3. pRPA2 foci alone do not predict overall survival.** Kaplan Meier curves evaluating overall survival between patients with pRPA2-High and pRPA2-Low tumors in the discovery cohort (n=31, HR 0.6 95% CI 0.2-1.5, *P*=0.3).

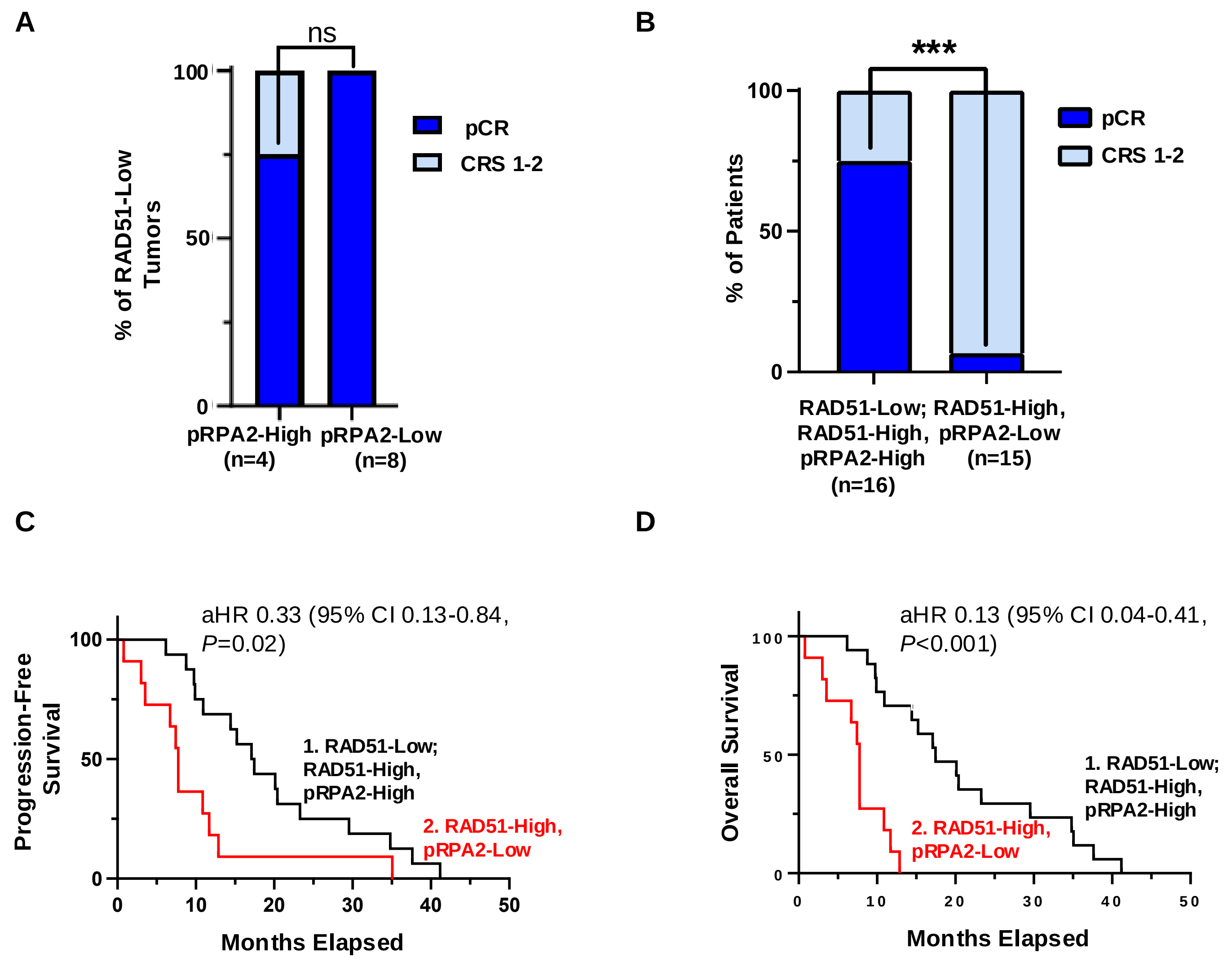


**No. at Risk**

| | | | | | | |
|---|---|---|---|---|---|---|
| 1. | 16 | 12 | 7 | 3 | 1 | 0 |
| 2. | 11 | 4 | 1 | 1 | 0 | 0 |

**No. at Risk**

| | | | | | | |
|---|---|---|---|---|---|---|
| 1. | 16 | 16 | 10 | 7 | 2 | 0 |
| 2. | 15 | 9 | 4 | 0 | 0 | 0 |

**Supplementary Figure 4. RAD51-Low and RAD51-High, pRPA2-High tumors are more sensitive to platinum chemotherapy than RAD51-High, pRPA2-Low tumors. A,** Proportion of patients with complete pathologic response (pCR) vs chemotherapy response score (CRS) of 1-2 in RAD51-Low tumors with either pRPA2-High or pRPA2-Low scores $P$=0.52. **B,** pCR in RAD51-Low or RAD51-High, pRPA2-High tumors vs RAD51-High, pRPA2-Low tumors $P$<0.001. **C,** Kaplan Meier curves evaluating progression-free survival (n=27, aHR 0.33, 95% CI 0.13-0.84, $P$<0.001) and **D,** Overall survival in patients stratified by RAD51/pRPA2 scores (n=31, aHR 0.13, 95% CI 0.04-0.41, $P$<0.001). *$P$<0.05, ***$P$<0.001 by student's two-tailed t-test. *aHR= adjusted hazard ratio for: age, stage, residual disease and *BRCA* status.

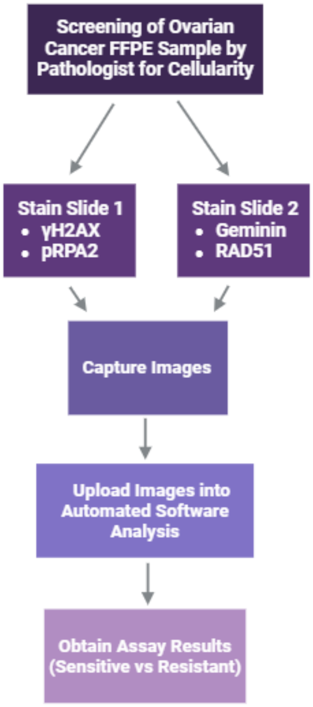
Screening of Ovarian Cancer FFPE Sample by Pathologist for Cellularity
Stain Slide 1
• γH2AX
• pRPA2
Stain Slide 2
• Geminin
• RAD51
Capture Images
Upload Images into Automated Software Analysis
Obtain Assay Results (Sensitive vs Resistant)

**Supplementary Figure 5. Schematic of functional RAD51 and pRPA2 foci assay.** Two FFPE slides are used, one is stained for pRPA2 and γH2AX and the other is stained for RAD51 and geminin. After imaging, images are uploaded to automated quantification software to obtain a score which is used to predict sensitivity or resistance.

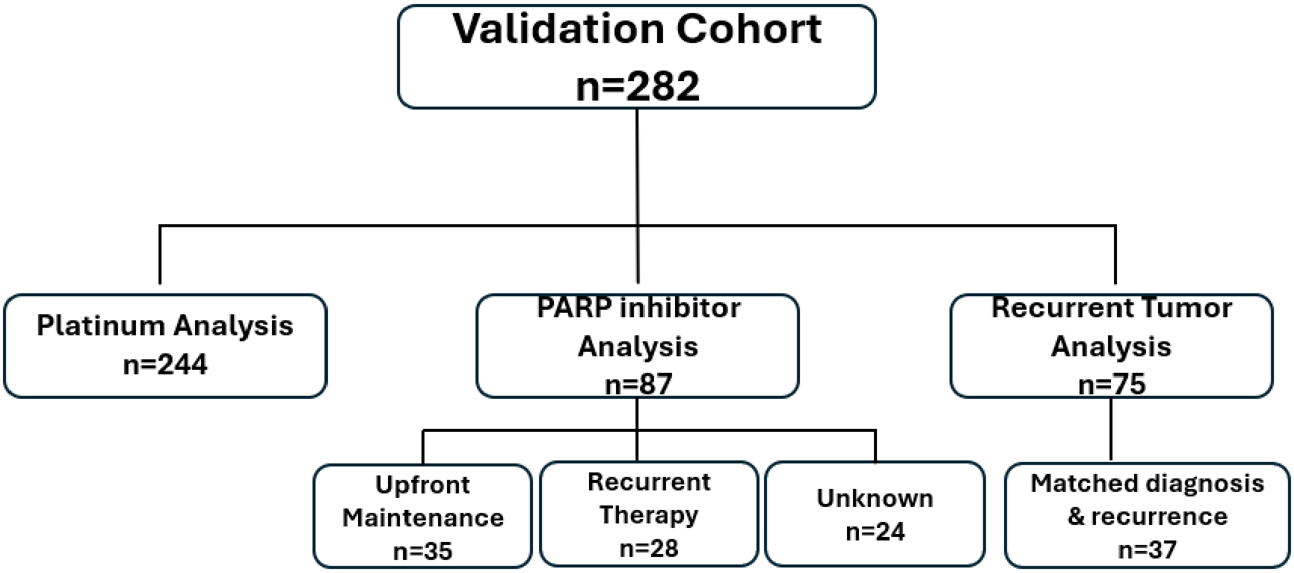
Validation Cohort
n=282
Platinum Analysis
n=244
PARP inhibitor
Analysis
n=87
Recurrent Tumor
Analysis
n=75
Upfront
Maintenance
n=35
Recurrent
Therapy
n=28
Unknown
n=24
Matched diagnosis
& recurrence
n=37

**Supplementary Figure 6. Flow diagram of high-grade serous ovarian cancer patient-derived samples used for the RAD51 and pRPA2 foci assay.** The validation cohort consisted of 244 ovarian cancer samples from patients who received platinum chemotherapy. Within this cohort, there was a subset of patients who received PAPR inhibitors (n=87). These tumors were analyzed to understand the role of the RAD51 and pRPA2 foci assay to predict PARP inhibitor response. Lastly, there was as subset of tumors obtained at the time of recurrence (n=38). Of these, 37 had had matched samples from diagnosis to recurrence. These samples were analyzed to understand the role of the RAD51 and pRPA2 foci assay in the recurrent setting.

**A**

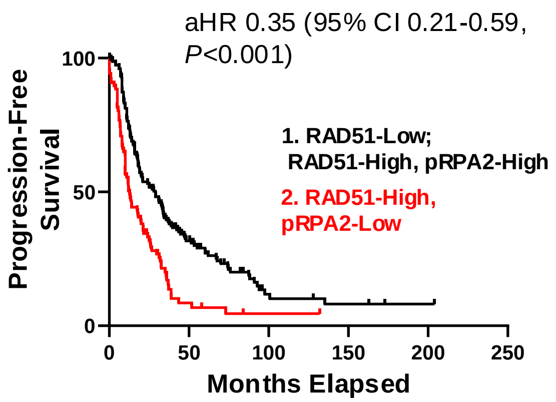


**No. at Risk**

| | | | | | | |
|---|---|---|---|---|---|---|
| 1. | 151 | 26 | 5 | 4 | 2 | 0 |
| 2. | 87 | 5 | 1 | 0 | 0 | 0 |

**B**

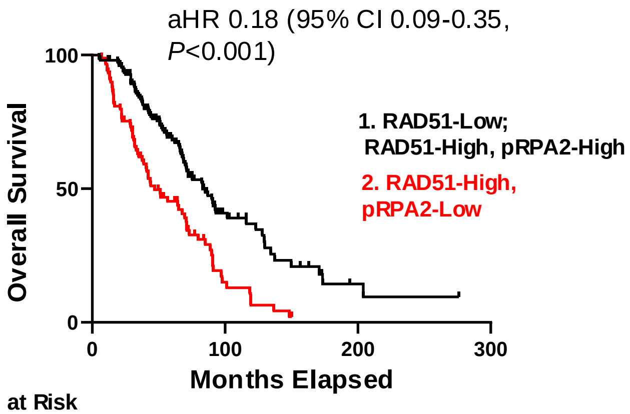


**No. at Risk**

| | | | | |
|---|---|---|---|---|
| 1. | 154 | 22 | 3 | 0 |
| 2. | 89 | 7 | 0 | 0 |

**Supplementary Figure 7. Combined RAD51 and pRPA2 scores predict survival in a validation cohort. A,** Kaplan Meier curves evaluating progression-free survival between patients with RAD51-Low or RAD51-High, pRPA2-High and RAD51-High, pRPA2-Low tumors (n=238, aHR 0.35, 95% CI 0.21-0.59 $P<0.001$) **B,** Kaplan Meier curves evaluating overall survival between patients with RAD51-Low or RAD51-High, pRPA2-High and RAD51-High, pRPA2-Low tumors (n=243, aHR 0.18, 95% CI 0.09-0.35 $P<0.001$). *aHR= adjusted hazard ratio for: age, stage, residual disease and *BRCA* status.

**A**

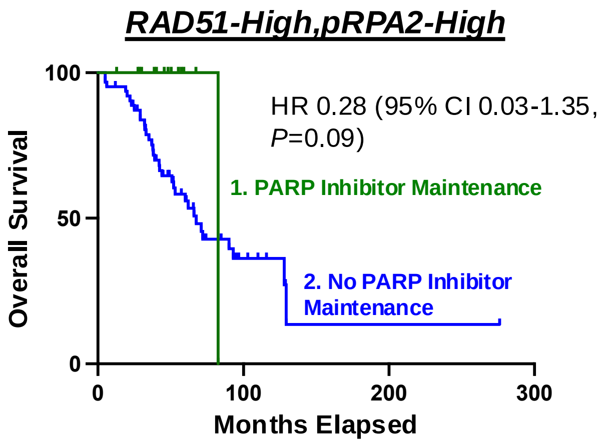


**No. at Risk**

| | | | | |
|---|---|---|---|---|
| 1. | 14 | 0 | 0 | 0 |
| 2. | 63 | 7 | 1 | 0 |

**B**

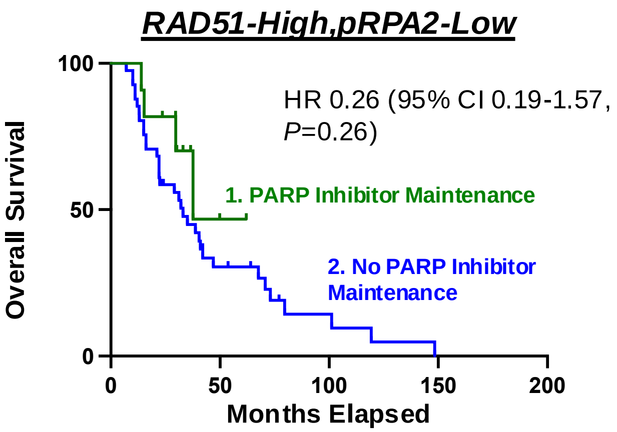


**No. at Risk**

| | | | | | |
|---|---|---|---|---|---|
| 1. | 41 | 10 | 3 | 0 | 0 |
| 2. | 11 | 1 | 0 | 0 | 0 |

**Supplementary Figure 8. Automated pRPA2 and RAD51 scores predict overall survival in patients with high grade serous ovarian cancer (HGSOC) receiving PARP inhibitors. A,** Kaplan Meier curves evaluating overall survival in patients with RAD51-High, pRPA2-High scores stratified by whether they received upfront PARP inhibitor maintenance therapy (n=77, aHR 0.28, 95% CI 0.03-1.35, $P$=0.09). **B,** Kaplan Meier curves evaluating overall survival in patients with RAD51-High, pRPA2-Low scores stratified by whether they received upfront PARP inhibitor maintenance therapy (n=52, aHR 0.26, 95% CI 0.19-1.57, $P$=0.26).
